# Activation of cannabinoid CB2 receptors by (−)-cannabichromene but not (+)-cannabichromene

**DOI:** 10.1101/2023.08.01.551199

**Authors:** Michael Udoh, Marina Santiago, Syed Haneef, Alison Rodger, Charles K. Marlowe, Philip J. Barr, Mark Connor

**Affiliations:** Macquarie Medical School, Macquarie University, NSW 2109, Australia; School of Natural Sciences, Macquarie University, NSW 2109, Australia; BayMedica, LLC, a Division of InMed Pharmaceuticals, 458 Carlton Court, South San Francisco, CA94080, U.S.A

## Abstract

**Introduction:** Cannabichromene (CBC) is a minor constituent of Cannabis that is a selective cannabinoid CB2 receptor agonist and activator of TRPA1. To date, it has not been shown whether (−)-CBC, (+)-CBC (or both) can mediate these effects. In this study we investigate the activity of the CBC enantiomers at CB1, CB2 and TRPA1 *in vitro*.

**Materials and Methods:** CBC enantiomers were purified from synthetic CBC by chiral chromatography, and their optical activity was confirmed by spectroscopy. Human CB1 and CB2 receptor activity was measured using a fluorescent assay of membrane potential in stably transfected AtT20 cells. TRPA1 activation was measured using a fluorescent assay of intracellular calcium in stably transfected HEK293 cells.

**Results:** (−)-CBC activated CB2 with an EC_50_ of 1.5 µM, to a maximum of 60 % of CP55940. (+)-CBC did not activate CB2 at concentrations up to 30 µM. Only 30 µM (−)– CBC produced detectable activation of CB1, (+)-CBC was inactive. Both (−)-CBC and (+)– CBC activated TRPA1; at 30 µM (−)-CBC produced an activation 50% of that of the reference agonist cinnamaldehyde (300 µM), 30 µM (+)-CBC activated TRPA1 to 38% of the cinnamaldehyde maximum.

**Discussion:** It is unclear whether (−)-CBC is the sole or even the predominant enantiomer of CBC enzymatically synthesized in *Cannabis*. This study shows that (−)-CBC is the active isomer at CB2 receptors, while both isomers activate TRPA1. The results suggest that medicinal preparations of CBC that target cannabinoid receptors would be most effective when (−)-CBC is the dominant isomer.

## Introduction

Cannabichromene (CBC) is one of the four most abundant phytocannabinoids in Cannabis (Izzo et al., 2009) and has recently been reported to be a selective cannabinoid CB2 receptor agonist, with an efficacy higher than that of the principal biologically active phytocannabinoid, Δ^9^-tetrahydrocannabinol (THC) (Udoh et al, 2019). CBC also activates the sensory stimulus-gated ion channel TRPA1 (de Petrocellis et al., 2008). Preclinical studies in mice suggest that CBC is anticonvulsant in a model of Dravet syndrome, (Anderson et al., 2021) and anti-inflammatory following chemically induced intestinal damage (Izzo et al., 2012, Romano et al., 2013). Cannabichromene is not regulated under United Nations Single Convention on Narcotic Drugs (International Narcotic Control Board, 2022), and CBC is available for human consumption in many parts of the United States and Canada; it is thus important to understand the structural basis of its biological activity.

Many phytocannabinoids are chiral (Filer, 2021) and the (−) isomer is usually the only one present in the plant, while synthetic procedures often produce racemic mixtures. CBC may be an exception to this with several papers reporting that CBC isolated from *Cannabis* consists of scalemic (unequal) mixtures of (−) and (+) CBC, (Agua et al, 2021, Calcaterra et al, 2023 Mazzoccanti et al 2017), and original studies suggested that natural CBC was a racemic mixture (Gaoni & Mechoulam, 1971). The production of (+) and (−) isomers of biologically important molecules in plants is not uncommon (Bitchagno et al, 2022). Intriguingly, the enzyme responsible for the synthesis of CBCA (the precursor of CBC) appears to preferentially (but not exclusively) produce (−)-CBCA (Morimoto et al., 1997). Understanding whether the biological activity is mediated by one isomer (or both) is important for understanding the effects of naturally derived and synthetic cannabinoids, as well as providing structural information that can aid in understanding receptor function and inform the design of new molecules. For example, in experiments where (−)-cannabidiol (CBD) does not affect binding of [^3^H]-HU243 to CB1 or CB2 receptors, (+)-CBD has a sub-micromolar affinity for both (Bisogno et al., 2001). Our previous study identifying the CB2 activity of CBC used synthetic (racemic) CBC. In this study we have examined the effects of resolved isomers of CBC on human CB2 receptors and TRPA1 channels. We find that the CB2 activity resides exclusively in the (−) isomer, while the activity of the (−) and (+) isomer at TRPA1 were similar.

## Methods

Details of cell culture conditions, media and solutions are described in the Supplementary material. All experiments were repeated at least six times, in duplicate, unless otherwise stated.

### CBC synthesis and purification

Racemic CBC was chemically synthesized at multi-gram scale using a slightly modified method of Agua et al. (2021). Three grams of the partially purified racemic mixture of CBC was applied to a chiral preparative HPLC column (Amylose AD-H, Regis Technologies, Inc., Illinois; mobile phase IPA-hexane) to achieve baseline separation. Individual chiral isomers were further purified at ∼100 mg scale by preparative thin layer chromatography (TLC) on silica gel (2 mm thickness) using 10% hexane/ethyl acetate as the mobile phase, to give clear colorless oils.

Following preparative TLC, the purity of each sample was verified by analytical chiral column HPLC (Lux® Amylose, Phenomenex, Supplementary Figure 1), and their structures were also verified by NMR and mass spectrometry. Dried material was dissolved in DMSO, coded and shipped to Macquarie University. The absorbance and circular dichroism (CD) spectra of the samples were determined on a Jasco (Hachioji, Japan) J-1500 spectropolarimeter by diluting the enantiomerically pure sample to 1 mg/ml in DMSO/ethanol. The slower eluting isomer showed a negative CD spectrum with a maximum at 276 nm; this was the same shape but the opposite sign from the CD spectrum of the faster eluting isomer (Supplementary Figure 2). The absorbance spectra do not overlay at shorter wavelengths suggesting that the faster eluting sample includes a likely achiral impurity. The coding was broken only after the *in vitro* experiments were completed. (−)CP55940 was from Cayman Chemical (Ann Arbor, MI, # 90084). Compounds were aliquoted in DMSO at 30 mM and stored at –20 °C, a fresh aliquot of drug was used each day. The final DMSO concentration was 0.1% in all experiments., except for the spectroscopy which was undertaken in 5% DMSO, resulting in a wavelength cutoff of 235 nm.

### Assay of CB receptor function

Changes in membrane potential were measured using a proprietary membrane potential assay kit (MPA Blue, #8034, Molecular Devices, Sunnyvale CA) as described previously (Knapman et al, 2013). AtT20 FlpIn cells stably expressing human CB1 and CB2 (AtT20-CB1, –CB2) were cultured overnight in L-15 media (supplemented with 1% FBS, 1 % penicillin/streptomycin and 15 mM glucose) and then incubated in a Flexstation 3 for at least 60 minutes at 37 °C with MPA dye dissolved in a modified Hanks Buffered Saline Solution (HBSS). Compounds were added in a volume of 20 µL (for a final volume of 200 µL) after a baseline recording of 60-120s. Readings of fluorescence were made every 2s; cells were illuminated at a wavelength of 530 nm, fluorescence was measured at 565 nm. Effects are reported as the peak change in fluorescence following compound addition, expressed as a percentage of the baseline. Values were corrected for the effects of the solvent (0.1% DMSO final concentration for all compounds).

### Assay of TRPA1 function

Changes in [Ca]_i_ were measured using a proprietary assay kit (FLIPR^®^ Ca5, #8186, Molecular Devices), as described previously (Redmond et al., 2014). HEK293 FlpIn/T-REx cells expressing human TRPA1 (HEK293-TRPA1) were cultured overnight in supplemented L-15 media and channel expression was induced 4 hours before drug addition by addition with of 1 µg ml^-1^ tetracycline. Cells were incubated in a Flexstation 3 for at least 60 minutes at 37°C in Ca5 dye plus 1.25 mM probenecid (Biotium #50027, Jomar Life Research, Australia) prior to the start of recording. Readings of fluorescence were made every 2s; cells were excited at a wavelength of 485 nm, fluorescence was measured at 525 nm. Compound effects are reported as the peak change in fluorescence following addition, expressed as a percentage of the baseline. Values were corrected for the effects of the solvent (0.1% DMSO final concentration for all drugs).

### Molecular docking

Molecular docking analysis was performed using AutoDock version 4.2.6. (Morris et al., 2009). Preliminary steps such as receptor and ligand preparation were done using AutoDock Tools (ADT) version 1.5.6. The X-ray crystal structure of human Cannabinoid receptor 2 bound to the agonist AM-12033 (PDB ID: 6KPC; Hua et al., 2020) was used for docking. The (−)-CBC and (+)-CBC, 3D conformers of cannabichromene were obtained from PubChem database with compound ID 76971722 and 21668219 respectively. For further details, see Supplementary Methods.

## Results

CP55940 produced a decrease in fluorescence in AtT20-CB2 cells with a pEC50 of 7.2 ± 0.05, with a maximum change in fluorescence of 34 ± 1 %. (−)-CBC produced a decrease in fluorescence with a pEC50 of 5.8 ± 0.2, with a maximum change in fluorescence of 23 ± 2 %. (+)-CBC did not significantly affect the membrane potential of AtT20 CB2 cells. At the highest concentration of (+)-CBC tested (30 µM), the change in fluorescence was 3 ± 2 %, the apparent maximum change in fluorescence when 2 vehicle traces were subtracted from each other was 2 ± 2 % (Figure 1), indicating that the effects of (+)-CBC were within the baseline variability in the assay.

**Figure 1:**
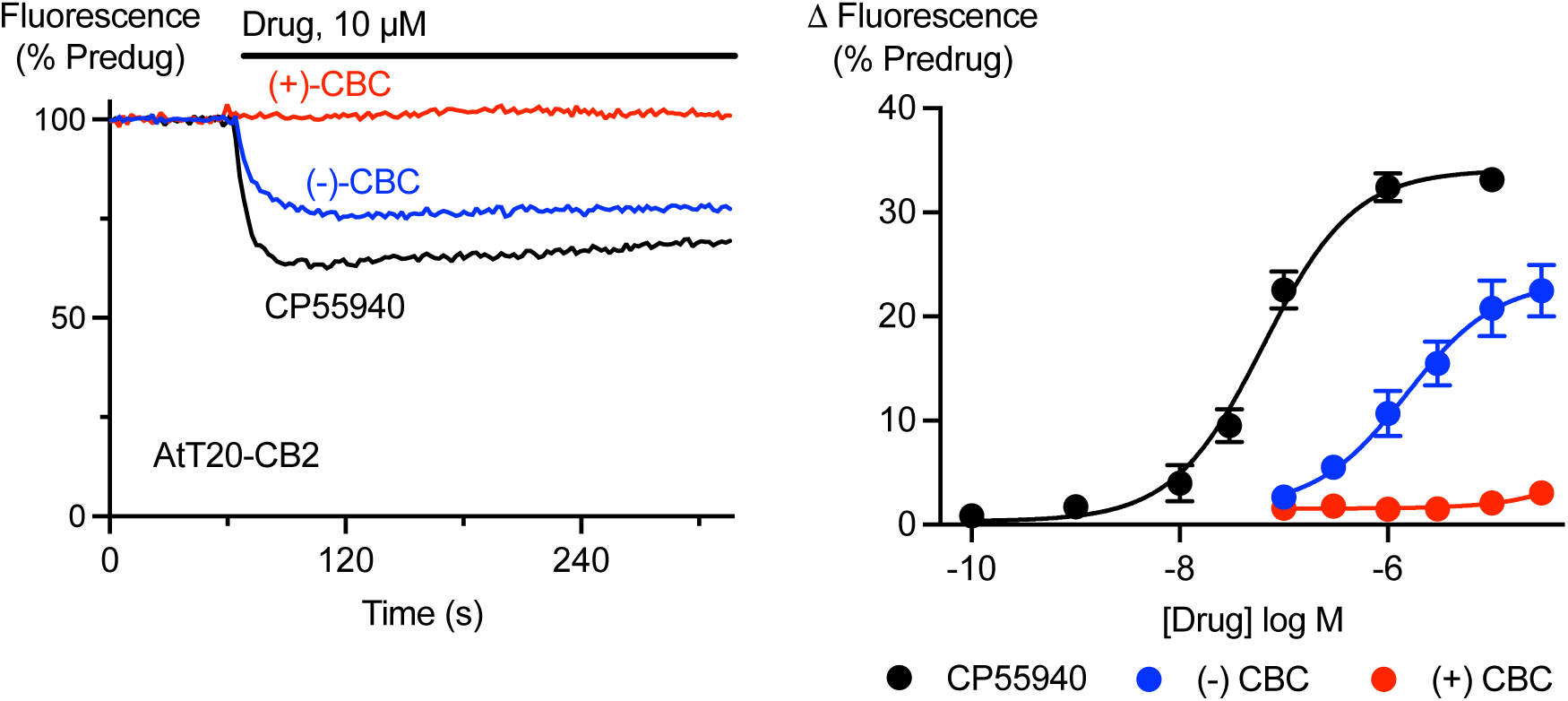
Activation of CB2 receptors by cannabichromene isomers. AtT20 cells expressing CB2 receptors were challenged with (−)-CBC, (+)-CBC and CP55940. A fluorescent membrane potential dye was used to measure the response. A) Representative traces showing that application of 10 µM (−)-CBC and CP55940 produced a sustained decrease in fluorescence, consistent with hyperpolarization of the cells, while 10 µM (+)-CBC had not effect. B) Concentration-response curves for (−)-CBC, (+)-CBC and CP55940 showing the maximum change in fluorescence produced by each drug. Each point represents the mean ± s.e.m. of at least 6 independent experiments, performed in duplicate.

In AtT20-CB1 cells, CP55940 produced a decrease in fluorescence with a pEC50 of 7.53 ± 0.06, with a maximum change in fluorescence of 33 ± 1 %. Only at the highest concentration tested (30 µM) did (−)-CBC produced a change in fluorescence (6 ± 1.4 %) that was significantly greater than that of vehicle (2.0 ± 0.6; P=0.0083; One-way ANOVA, P=0.034, followed by Dunnet’s multiple comparisons test, each (−)-CBC concentration compared with vehicle). For (+)-CBC (30 µM) the change in fluorescence was 2.2 ± 0.4 %, vehicle was 3.1 + 0.8 % (Figure 2).

**Figure 2:**
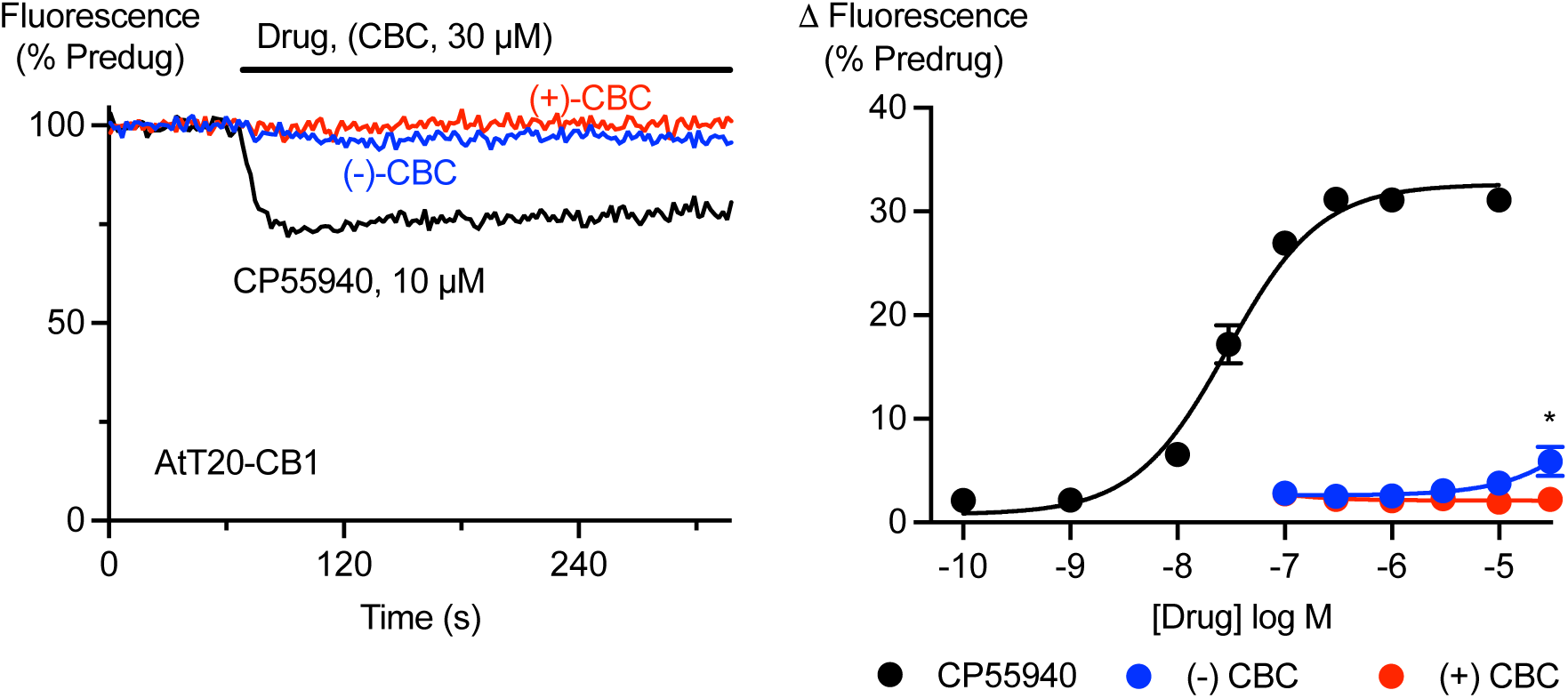
Activation of CB1 receptors by cannabichromene isomers. AtT20 cells expressing CB1 receptors were challenged with (−)-CBC, (+)-CBC and CP55940. A fluorescent membrane potential dye was used to measure the response. A) Representative traces showing that application of 10 µM CP55940 produced a sustained decrease in fluorescence, consistent with hyperpolarization of the cells, while 30 µM (−)-CBC had a minimal effect and 30 µM (+)-CBC had no effect. B) Concentration-response curves for (−)-CBC, (+)-CBC and CP55940 showing the maximum change in fluorescence produced by each drug. Each point represents the mean ± s.e.m. of at least 6 independent experiments, performed in duplicate. Only 30 µM (−)-CBC only produced a significant change in fluorescence (One-way ANOVA, P=0.034, followed by Dunnet’s multiple comparisons test, * P=0.0083; each concentration compared with vehicle).

In HEK293-TRPA1 cells cinnamaldehyde produced an increase in Ca5 fluorescence of 384 ± 36 %, with a *p*EC_50_ of 4.96 ± 0.06. Both isomers of CBC activated TRPA1. (−)-CBC appeared more potent than (+)-CBC, at 10 µM (−)-CBC produced an increase in fluorescence of 113 ± 27 %, while the increase produced by (+)-CBC was 29 ± 16 % (Figure 3). At the highest concentration tested, 30 µM, the increase in fluorescence produced by (−)-CBC was 202 ± 27 %, and the increase produced by (+)-CBC was 159 ± 27 %. It was not possible to estimate the maximum effect of either CBC isomer (Figure 3), and so the *p*EC_50_ could not be estimated.

**Figure 3:**
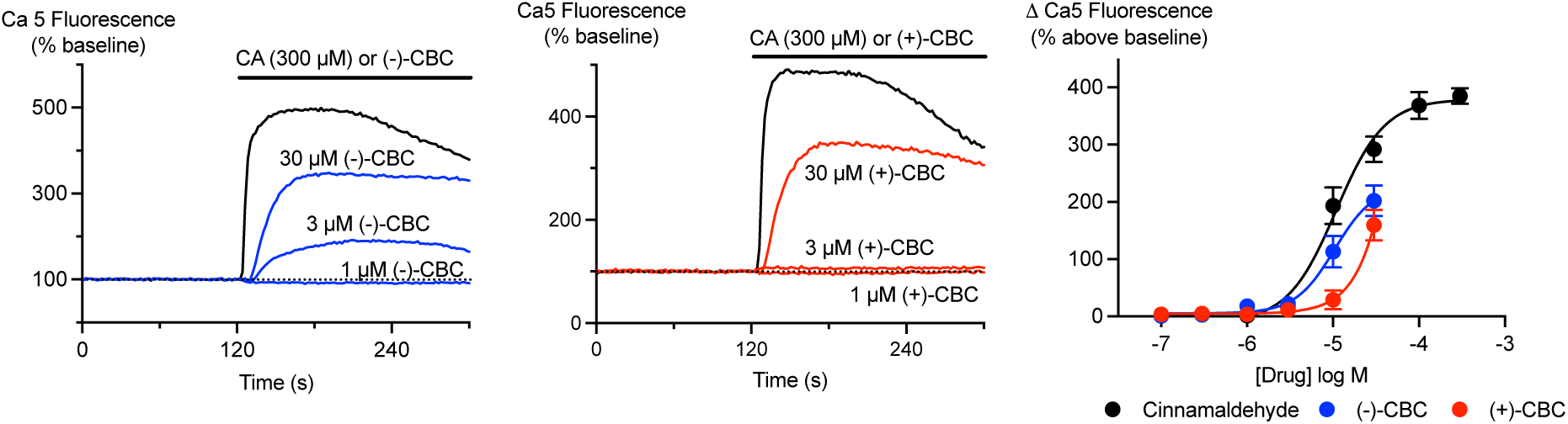
Activation of TRPA1 by cannabichromene isomers. HEK293 cells expressing TRPA1 receptors were challenged with (−)-CBC, (+)-CBC and cinnamaldehyde (CA). Ca5 dye was used to measure changes in intracellular Ca. A) Representative traces showing that application of 300 µM cinnamaldehyde (CA) produced a sustained increase in fluorescence, with smaller responses to 30 and 3 µM (−)-CBC. B) Representative traces from cells where (+)-CBC was applied, again 30 µM produced a smaller change in intracellular Ca than CA, and 3 µM (+)-CBC was largely without effect. C) Concentration-response curves for (−)-CBC, (+)-CBC and CA showing the maximum change in fluorescence produced by each drug. Each point represents the mean ± s.e.m. of at least 6 independent experiments, performed in duplicate.

In an earlier set of experiments we examined the effects of less well-resolved preparations of the isomers (see Methods); in these experiments the preparations contained of 97 % “fast” or “slow” (referring to their relative HPLC mobilities) isomer and 3 % of the other isomer (Figure 4). In AtT20-CB2 cells, the mixture containing predominantly (+)-CBC (102D) produced a modest hyperpolarization, with a maximum change in fluorescence of 13 ± 4.3 % at 30 µM. By contrast, the mixture containing predominantly (−)-CBC (102E) produced a hyperpolarization with a *p*EC_50_ of 5.86 ± 0.11, to a maximum of 26 ± 2 % at 30 µM. Synthetic (+/-)-CBC hyperpolarized AtT20-CB2 cells with a *p*EC_50_ of 5.56 ± 0.12, to a maximum of 24 ± 2% at 30 µM. CP55940 hyperpolarized AtT20-CB2 cells with a *p*EC_50_ of 7.5 ± 0.1, with a maximum change in fluorescence of 35 ± 2 %. The EC_50_ of 102E (1.39 µM) is about half that of the (+/-)-CBC (2.76 µM), providing further evidence that the (−)-CBC is responsible for most of the activity of synthetic CBC at CB2. The hyperpolarization produced by 30 µM 102D (13 ± 4 %) is similar to that produced by 1 µM 102E (13 ± 1%), suggesting that the activity in the 102D mixture arises from the presence of 3% (−)-CBC. The stereochemical dependence of CBC activation of TRPA1 is less pronounced than for CB2; the effects of (+/-)-CBC, 102D and 102E at TRPA1 are illustrated in Supplementary Figure 3.

**Figure 4:**
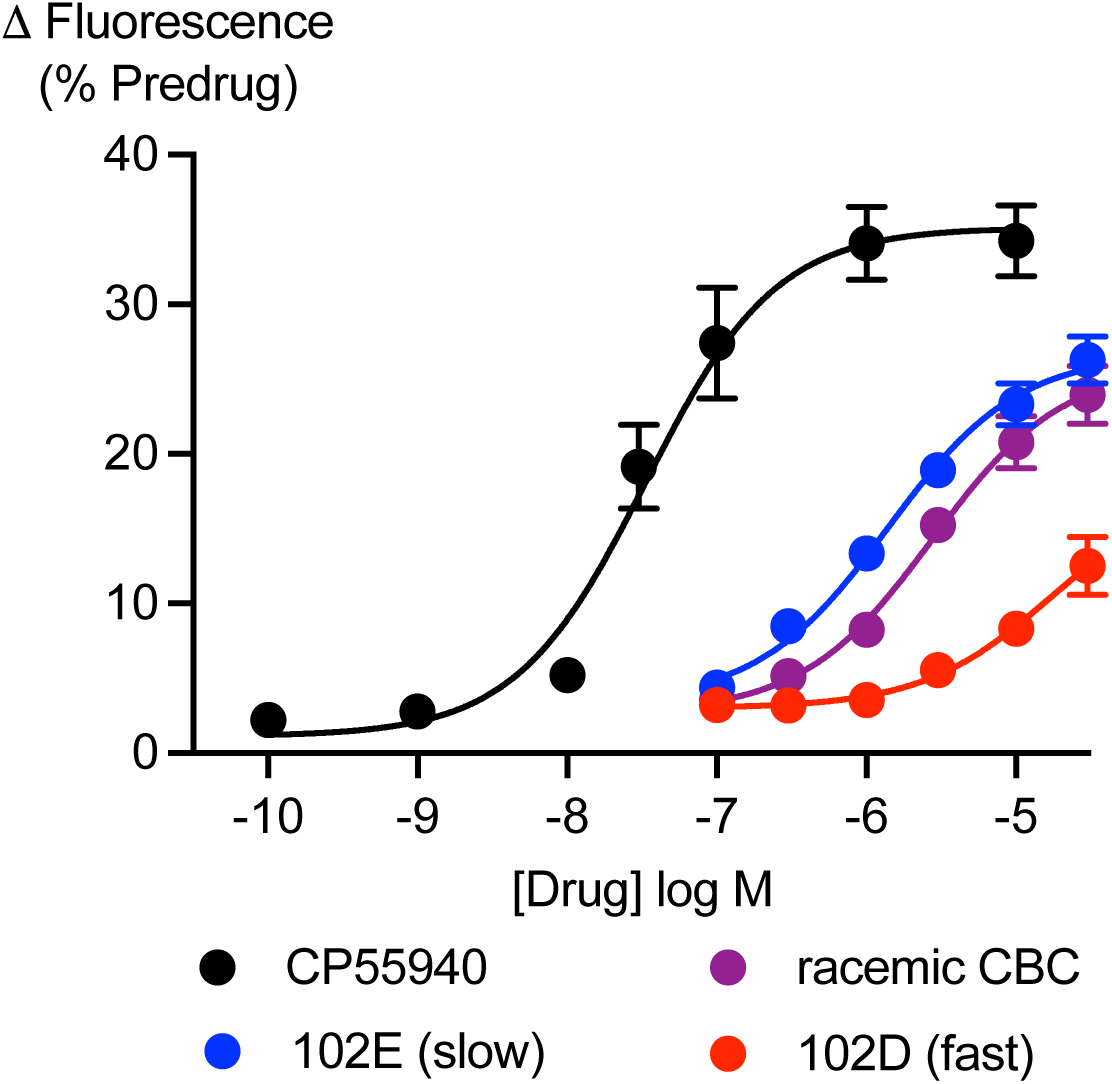
Activation of CB2 receptors by racemic or scalemic mixtures of CBC isomers. AtT20 cells expressing CB2 receptors were challenged with CP55940, (+/-)-CBC, 102E (97% (−), 3 % (+)-CBC) and 102D (97% (+) and 3 % (−) CBC). A fluorescent membrane potential dye was used to measure the response. Concentration-response curves for each drug or mixture showing the maximum change in fluorescence produced by each drug. Each point represents the mean ± s.e.m. of 5 independent experiments, performed in duplicate.

The isomers of CBC were docked with the computed structure of agonist-bound CB2 and were computed to interact with a mostly similar set of amino acids (Supplementary Figures 4, 5). For (−)-CBC, these included Ile110, Phe94 and Phe83, all of which were identified as forming part of the CB2 binding pocket accessed by the tetrahydrocannabinol-derivative AM12033 (Hua et al; 2020). Both (−)-CBC and (+)-CBC also appeared to interact with His95 and Val13, additional residues identified as part of the CB2 binding pocket of the aminoalkylindole CB agonist WIN55212 (Xing et al., 2020). (+)-CBC did not apparently interact with Phe94, but the binding energy (enthalpy) calculated for both isomers was similar (−11.78 Kcal/mol for (−)-CBC; –11.51 Kcal/mol for (+)-CBC). No attempt was made to estimate the entropic contribution to binding.

## Discussion

The principal finding of this study is that (−)-CBC has much higher activity at CB2 than does (+)-CBC. Further, although both enantiomers were able to activate TRPA1, (−)-CBC appeared to be more potent, although it was not possible to complete full concentration-response curves. The results confirm our previous results using synthetic CBC from a different source demonstrating that CBC is a CB2 selective agonist (Udoh et al, 2019). A preliminary analysis of the binding of (−) and (+)-CBC to the human CB2 receptor using computational methods found little difference between the free energy of binding for the molecules, although (−)-CBC appeared to interact with several additional residues of the receptor compared with (+)-CBC, including Phe94, a residue forming part of the binding pocket of the CB2 agonists AM12033 and WIN55212.

We have consistently found that CBC is a CB2-preferring agonist, although another study (Zazoog et al, 2020) found some activity of CBC in an assay of cAMP accumulation in both CB1 and CB2 transfected CHO cells. The reason for this inconsistency is not clear. The assay we used, activation of G protein-gated K channels, is not subject to strong amplification, and requires significant receptor occupancy in AtT20 cells. Nevertheless, in this assay the low efficacy phytocannabinoid THC produces a maximum effect between 50% and 75% of that of high-efficacy agonists such as CP55940 or 2-AG at CB1 receptors (Sachdev et al., 2019, Udoh et al., 2019). We did see a very small effect of (−)-CBC (30 µM) in AtT20-CB1 cells; this low efficacy could be amplified by the assay conditions and receptor expression levels in the study of Zazoog and colleagues (2020). In a study of autaptic hippocampal neuron neurotransmission, which is regulated by CB1 but not CB2 receptors, CBC was inactive (Straiker et al., 2021).

The potency and apparent efficacy of CBC to activate TRPA1 is also much less than reported in the study of de Petrocellis and colleagues (2001) on recombinant rat TRPA1. Again, the reasons for the differences have not been established, but are likely due to assay conditions and potentially species differences. de Petrocellis et al. (2001) measured TRPA1 activation following Ca addition to a cell suspension of HEK293 cells expressing TRPA1, while we used adherent HEK293 cells expressing human TRPA1. It is possible that responses at TRPA1 are sensitized in cells subject to the mechanical forces of being stirred suspension following exposure to a protease. Interestingly, de Petrocellis and colleagues (2001) also measured CBC elevations of Ca in adherent neonatal sensory neurons and CBC potency in these cells was 34 µM, which is in the same range as observed in our study.

The (−) isomers of phytocannabinoids have higher efficacy at CB receptors, (Edery et al., 1970) with the notable exception of (+)-CBD, which has a higher affinity for CB1 and CB2 than the naturally occurring (−)-CBD (Bisogno et al; 2001). Cannabis appears to contain racemic or scalemic mixtures of CBC (Gaoni & Mechoulam, 1971; Mazzoccanti et al, 2017; Agua et al., 2021, Calcaterra et al 2023) but, based on our findings, Cannabis extracts and synthetic products with a predominance of (−)-CBC are expected to have greater potential for CB2-mediated therapeutic effects than scalemic mixtures or preparations of (+)-CBC.

## Acknowledgements

Michael Udoh and Syed Haneef Syed Askar were supported by International Research Excellence Scholarships from Macquarie University. The CD was supported by the Australian Research Council Industrial Transformation Centre for Facilitated Advancement of Australia’s Bioactives (Grant IC210100040) and Research Attraction and Acceleration Program funding from the Office of the Chief Scientist and Engineer, Investment NSW. MU, MS, MC collected and analyzed biological data; PJB, CKM purified and characterised the CBC isomers, AR did the CD, SH modelled. MC wrote the draft MS all authors contributed with additions and revisions.

---

**Table 1:** Responses of CBC preparations at human cannabinoid receptors and TRPA1.

| | CB2<br>EC50, Max* | CB1<br>Response at 30 μM* | TRPA1<br>Response at 30 μM** |
| --- | --- | --- | --- |
| (-)-CBC | 1.5 μM, 60 ± 6 % | 17 ± 4% | 50 ± 6 |
| (+)-CBC | n.d., 10 ± 3 % @ 30 μM | 6 ± 1 % | 38 ± 8 |
| “slow”-CBC | 1.4 μM, 70 ± 3 % | n.d. | 69 ± 6 |
| “fast” CBC | n.d., 37 ± 4 % @ 30 μM | n.d. | 56 ± 8 |
| (+/-)-CBC<br>(synthetic) | 2.8 μM, 70 ± 3 | n.d. | 74 ± 6 |

n.d. – not determined
\* % 1 μM CP55940
\*\* % 300 μM cinnamaldehyde

**Supplementary Figure 1:**
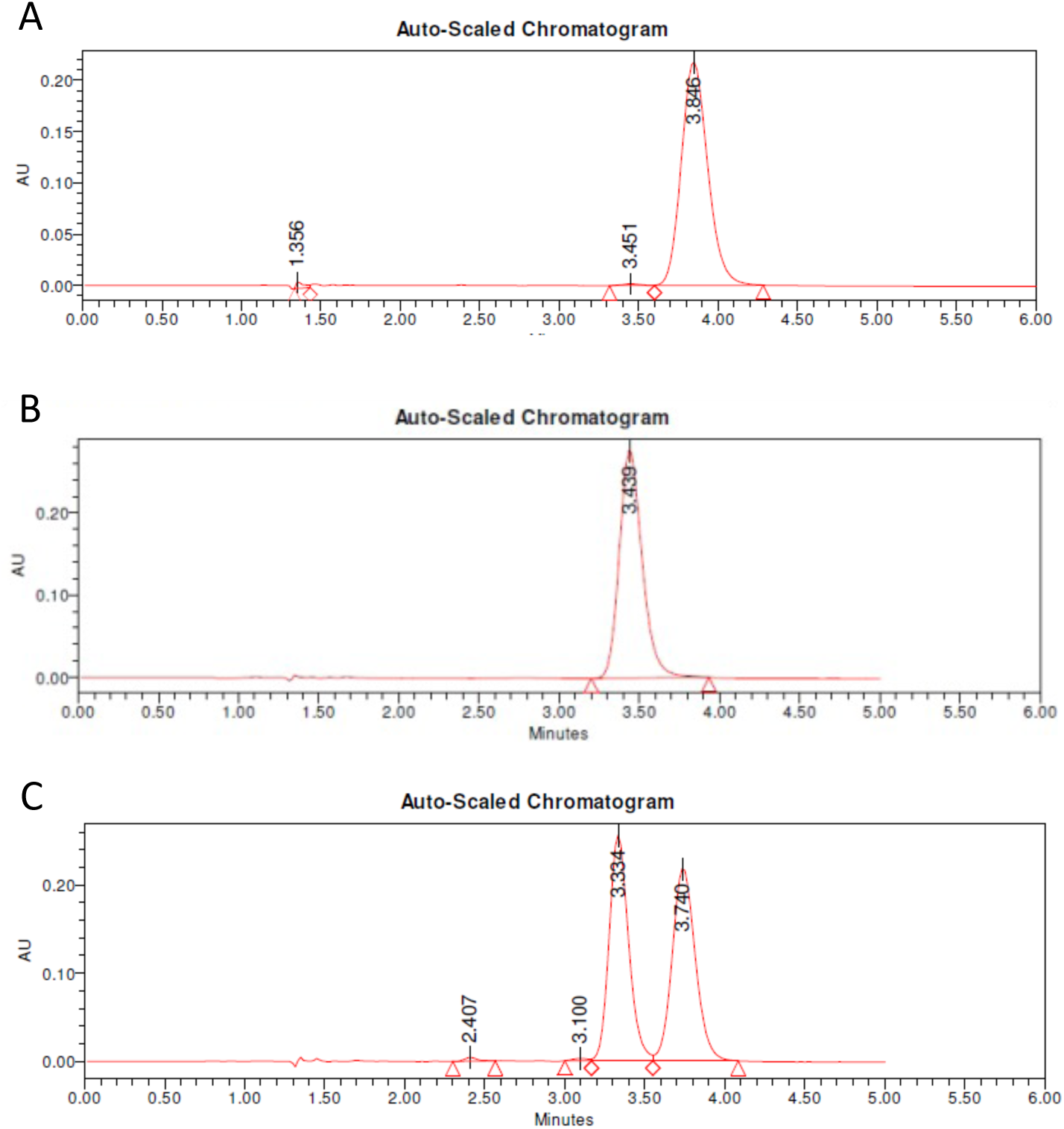
Chromatograms of purified CBC isomers and racemic CBC used in this study. Example chromatograms illustrating the retention times of A) purified (−)-CBC; B) purified (+)-CBC and synthetic, racemic CBC. Note the faster elution time of (+)-CBC in these conditions. The traces represent auto-scaled chromatograms after the injection of 5 µl of 1 mM sample on a Lux^®^ 5 µm Amylose-1 LC-column, 250 x 50 mm, AXIA™ Packed using an isocratic gradient of 20% water and 80% acetonitrile with a flow rate of 2ml/min.

**Supplementary Figure 2:**
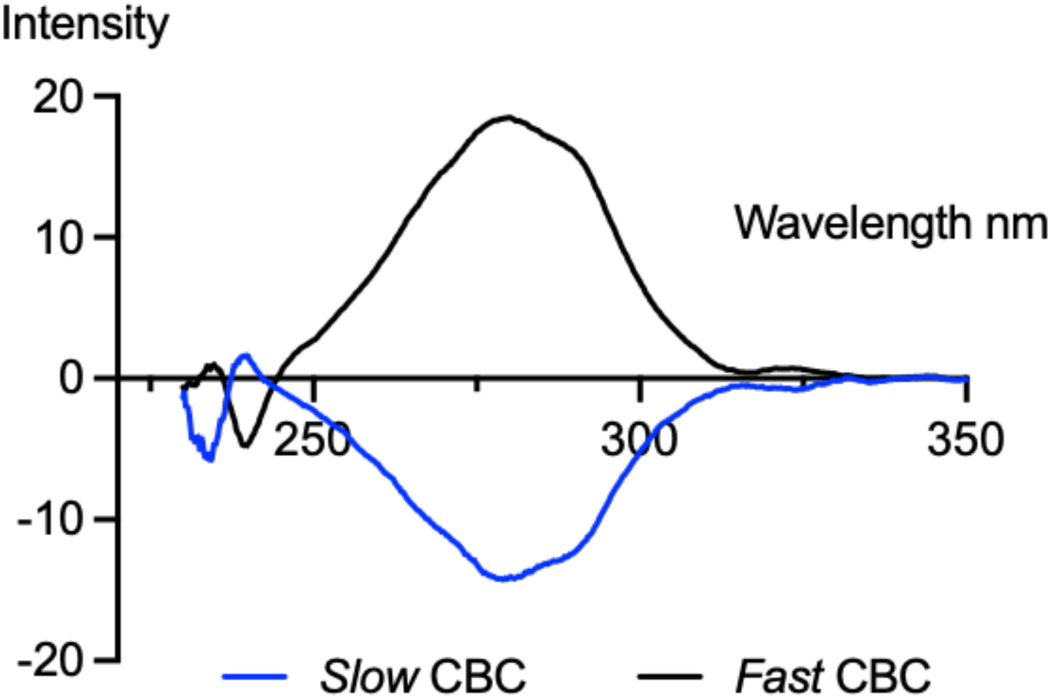
CD Spectrum of Fast and Slow CBC. The absorbance and circular dichroism (CD) spectra of the samples were determined on a Jasco (Hachioji, Japan) J-1500 spectropolarimeter by diluting the enantiomerically pure sample to 1 mg/ml in DMSO/ethanol, 1 mm path length.

**Supplementary Figure 3:**
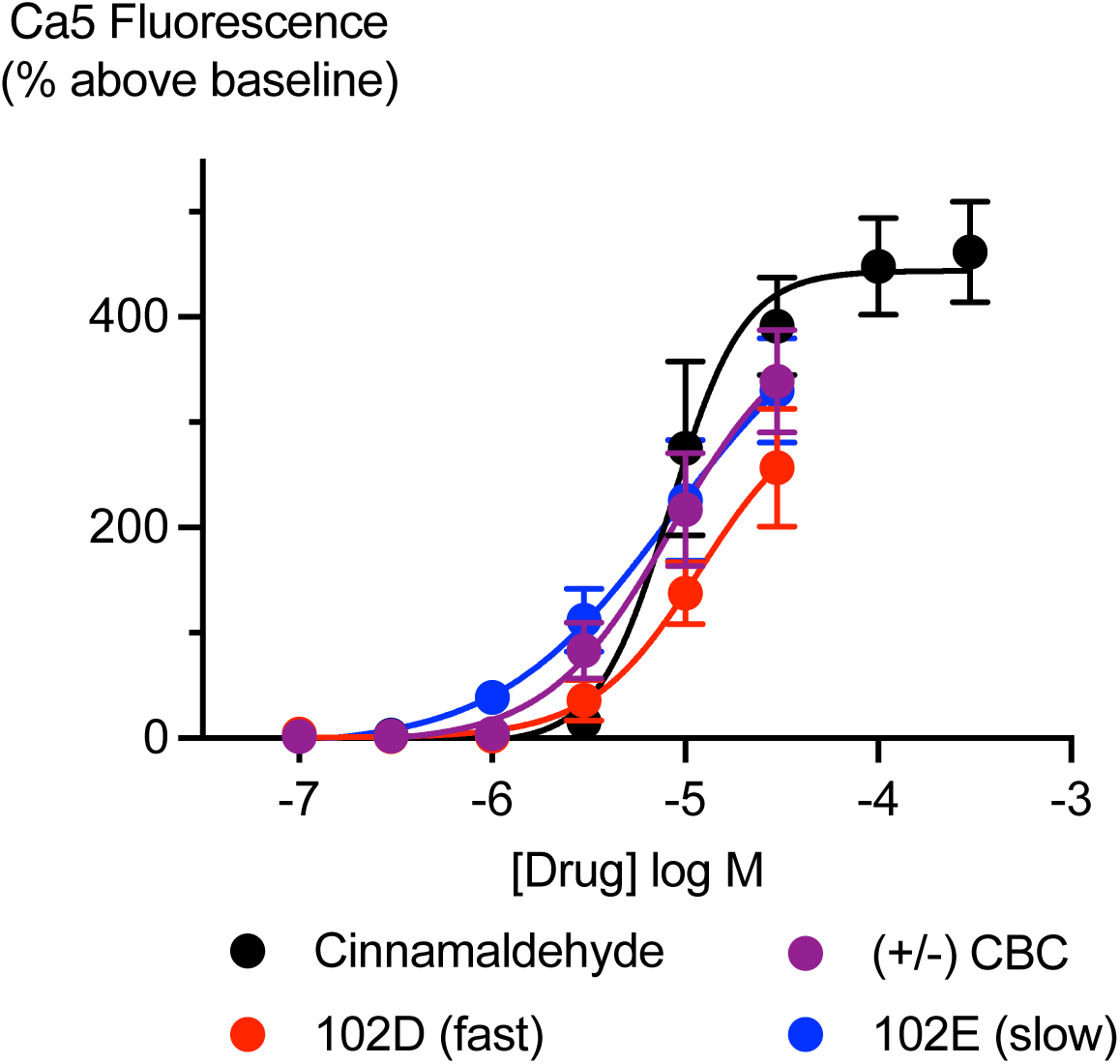
Activation of TRPA1 receptors by racemic or scalemic mixtures of CBC isomers. HEK293 cells expressing TRPA1 were challenged with cinnamaldehyde, (+/-)-CBC, 102E (97% (−), 3 % (+)-CBC) and 102D (97% (+) and 3 % (−) CBC). Ca5 dye was used to measure the response. Concentration-response curves for each drug or mixture showing the maximum change in fluorescence produced by each drug. Each point represents the mean ± s.e.m. of 5 independent experiments, performed in duplicate.

**Supplementary Figure 4:**
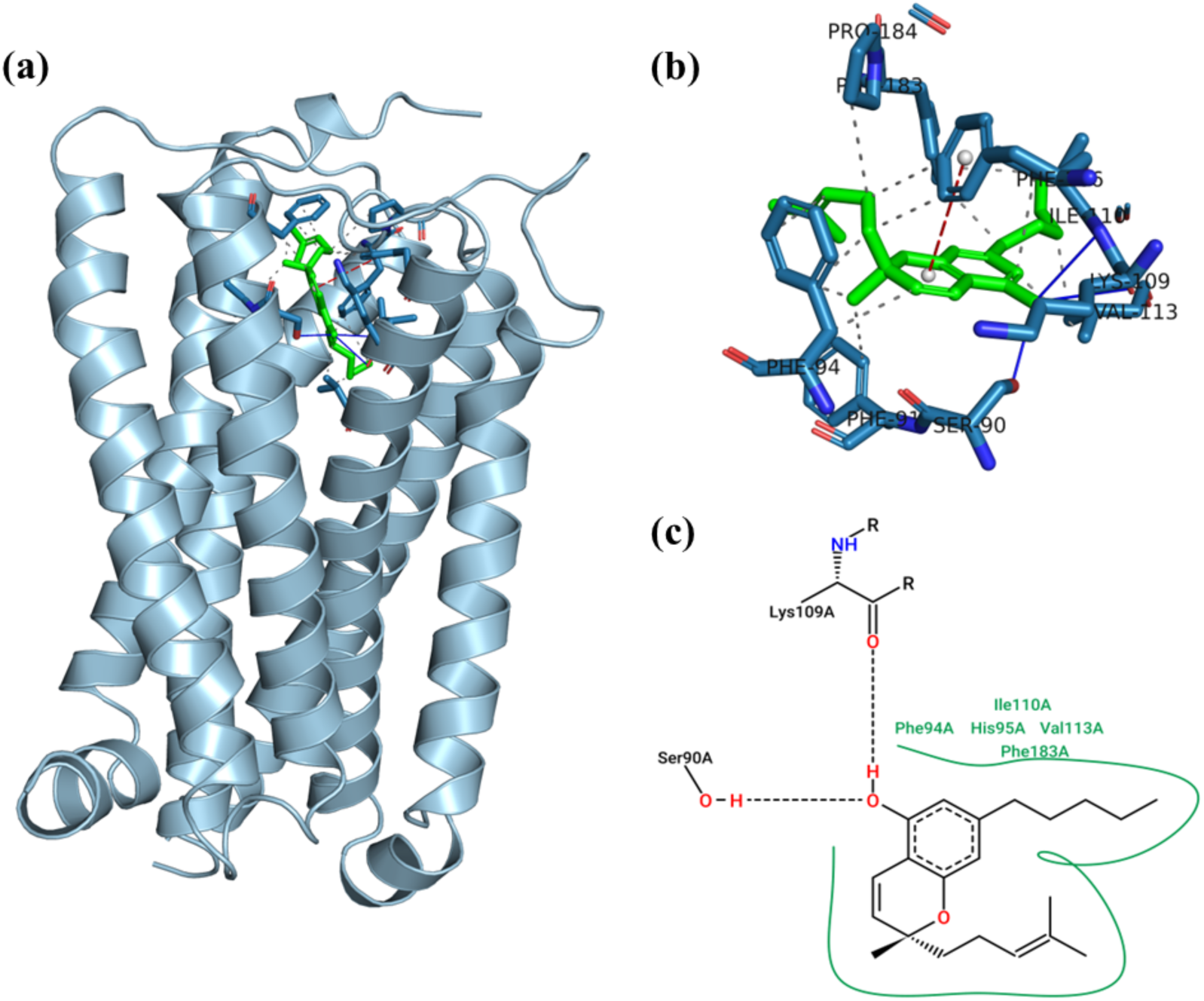
Visualization of (−)-CBC docked with the CB2 receptor. Cartoon representation of ligand-receptor docked complex, (b) Zoomed in version of ligand-receptor docked complex with (−) CBC isomer (*green colour, stick representation*) with interacting receptor residues, Hydrogen bond and Pi-Pi interactions shown in blue lines and red dash lines respectively and, (c) 2D PoseView of ligand-receptor interaction.

**Supplementary Figure 5:**
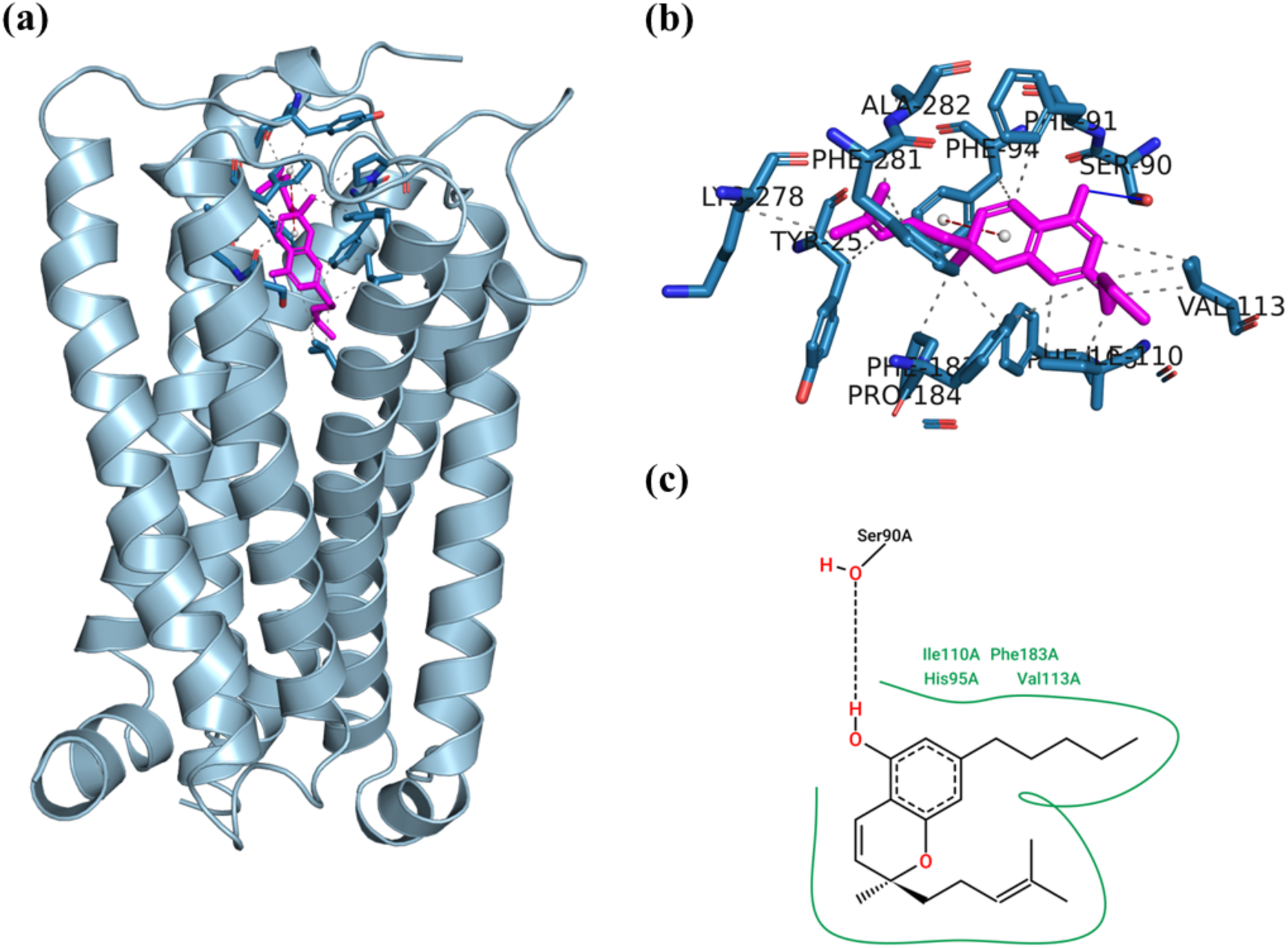
Visualization of (+)-CBC docked with the CB2 receptor. Cartoon representation of ligand-receptor docked complex, (b) Zoomed in version of ligand-receptor docked complex with (+) CBC isomer (*green colour, stick representation*) with interacting receptor residues, Hydrogen bond and Pi-Pi interactions shown in blue lines and red dash lines respectively and, (c) 2D PoseView of ligand-receptor interaction.

## Supplementary Information

*Cell culture:* AtT20FlpIn-CB1 and –CB2 cells were cultured in DMEM (GIBCO #11995-065) supplemented with 10 % FBS (Bovogen #SFBS, East Keilor, Victoria, Australia), 1% penicillin/streptomycin (GIBCO #15140122) 80 µg ml^-1^ hygromycin B gold (Invivogen#ant-hg). Cells were grown in 75 mm^2^ flasks (Falcon or Corning) at 37°C in a humidified incubator with 5% CO2. For experiments, cells from ∼ 90% confluent flasks were detached using trpysin/EDTA (Sigma-Aldrich, #T4049) and plated in a 96 well black walled plate (Corning 3603) in L-15 media (GIBCO #11415-064) supplemented with 1% FBS and 1% penicillin/streptomycin. For experiments, MPA and Ca5 dye were dissolved in HBSS of composition: (in mM): NaCl 145, HEPES 22, Na_2_HPO_4_ 0.338, NaHCO_3_ 4.17, KH_2_PO_4_ 0.441, MgSO_4_ 0.407, MgCl_2_ 0.493, CaCl_2_ 1.26, Glucose 5.56, pH 7.4, osmolarity 310-320. Both dyes were used at 50% of the manufacturers recommended concentration. HEK293FlpIn/TRex-TRPA1 cells were cultured as above, except the DMEM was supplemented with blasticidin 150 µg/ml (Invivogen #ant-bl) and hygromycin (Invivogen #ant-hg).

*Compounds:* Cannabinoids and analogs were diluted in HBSS supplemented with 0.1% bovine serum albumin (SIGMA #A7030) to minimize non-specific binding to plastic. A final concentration of 0.1% DMSO was maintained throughout. Compounds were added to the plates at 10x their final concentration by the Flexstation.

*Docking:* Molecular docking analysis was performed using AutoDock version 4.2.6. Preliminary steps were done using AutoDock Tools (ADT) version 1.5.6. The X-ray crystal structure of human Cannabinoid receptor 2 (PDB ID: 6KPC) was used for docking. The native ligand from the receptor was removed, and the structure was refined for missing atoms. Polar hydrogen atoms were added and the Kollman charges were assigned. AutoGrid was used to prepare the grid map, and to calculate the affinity, desolvation and electrostatic map of each atoms using a grid box. The grid size was set to 60 × 60 × 60 XYZ points with grid spacing of 0.375 Å and grid center was assigned at dimensions X: 9.05, Y: –1.00 and Z: –44.63. These grid center dimensions were based on the 6KPC structure. The CBC isomers were loaded, ligand root was detected and both isomers had 8 torsional degrees of free rotatable bonds. Ligand conformation search was based on Lamarckian genetic algorithm. Genetic algorithm (GA) was run for at least 50 sets with at least 500 as the population size. AutoDock was used for docking with the prepared receptor and ligand along with the grid parameters. The iterations below 1.0 Å in root-mean-square deviation (RMSD) were clustered and ranked based on their binding free energy. The ligand pose with lowest energy of binding or binding affinity was extracted. AutoDock inhibition constant (Ki) was noted for each isomer and finally the receptor-ligand complex was built and saved for further analysis. The docked complex was analysed using PLIP (Protein-Ligand Interaction Profiler) server. The 2D representation of receptor-ligand interaction was generated using PoseView from ProteinsPlus server. Other graphical representation of docked complex and binding mode figures were prepared using PyMol visualization software (http://www.pymol.org).

